# GABAergic signaling shapes multiple aspects of *Drosophila* courtship motor behavior

**DOI:** 10.1101/2023.01.24.525304

**Authors:** Hoger Amin, Stella S. Nolte, Bijayalaxmi Swain, Anne C. von Philipsborn

**Author notes:** equally shared authorship (Hoger Amin, Stella S. Nolte, Bijayalaxmi Swain).

## Abstract

Inhibitory neurons are essential for nervous system function. GABA is the most important neurotransmitter for fast neuronal inhibition in vertebrates and invertebrates. GABAergic signaling in sex specific, *fruitless* expressing neuronal circuits of *Drosophila* is required for multiple aspects of male courtship behavior. RNAi mediated knockdown of the GABA producing enzyme Gad1 and the ionotropic receptor Rdl in the ventral nerve cord causes uncoordinated and futile copulation attempts, defects in wing extension choice and severe alterations of courtship song. Altered song of GABA depleted males fails to stimulate female receptivity, but rescue of song patterning alone is not sufficient to rescue male mating success. Knockdown of Gad1 and Rdl in brain circuits abolishes courtship conditioning. We characterize the around 220 neurons coexpressing Gad1 and Fruitless in the *Drosophila* male nervous system and propose inhibitory circuit motifs underlying key features of courtship behavior based on the observed phenotypes.

## Introduction

Neuronal inhibition is a universal feature of nervous systems. Fast neuronal inhibition is accomplished by GABA, a neurotransmitter conserved across vertebrates and invertebrates that is already present in the simple nerve nets of cnidaria (Rentzsch et al. 2019, Arendt 2020, Pierobon 2021). GABAergic inhibition in the nervous system is important for stabilizing and shaping network activity and counterbalancing excitation, allowing for efficient coding and preventing runaway excitation and epileptic seizure (Cossart et al. 2005, Hennequin et al. 2017, Yu and Yu 2017, Smart and Stephenson 2019). Additional from this very global and general role, inhibitory connections shape circuit motifs that enable computations performed in many different pathways across many different species (Braganza and Beck 2018, Huang and Paul 2019). Such circuit motifs support effective sensory processing, and adaptive behavioral choice in response to external stimuli and internal state (Baca et al. 2008, Gaudry and Kristan 2009, Hangya et al. 2014, Swanson and Maffei 2019). For example, inhibitory interneuron networks enable male moths to track female pheromone plumes, crickets to process the temporal structure of the intraspecific acoustic calls and honeybees to decode the waggle dance vibrational pattern of their nest mates (Ai et al. 2018). In the *Drosophila* larvae nervous system, inhibitory circuit motifs based on reciprocal inhibition, lateral disinhibition and feedback disinhibition allow the animal to respond to mechanical stimuli with two distinct motor programs, either head turning or head retracting (Jovanovic et al. 2016).

Here, we use *Drosophila* male courtship behavior to address the question how neuronal inhibition shapes a more complex behavioral sequence. Courtship consist of multiple coordinated and ordered steps relying on multimodal sensory integration and the generation of specific, precisely timed motor patterns. Characteristic courtship displays are the male tracking and following the female, tapping her abdomen with the foreleg to sample pheromones and vibrating one extended wing. The wing vibrations produce a highly structured acoustic signal, the courtship song that stimulates the female’s receptivity (Ellendersen and von Philipsborn 2017, Swain and von Philipsborn 2020). When the female slows down, the male attempts copulation by probing the female genitalia with his proboscis, bending his abdomen and bringing his genitalia in apposition with the female genitalia. If the female opens her vaginal plates (hypogynia), copulation can occur (Mezzera et al. 2020, Wang et al. 2021).

The neuronal circuits underlying male fly courtship are among the best-studied model circuits for behavior in the *Drosophila* nervous system (Auer et al. 2016, Ellendersen and von Philipsborn 2017, Rings and Goodwin 2019, Peng et al. 2021). Most, if not all key neuronal components for male courtship express the male specific transcription factor FruitlessM (FruM). FruM is present in around 2% cells of the adult nervous system (Stockinger et al. 2005, Manoli et al. 2005). Among them are neurons controlling following (Ribeiro et al. 2018), singing (von Philipsborn et al. 2011) and copulation attempts (McKellar et al. 2019). Several studies have surveyed the distribution, anatomy and interconnection of FruM expressing neurons (Billeter and Goodwin 2004, Stockinger et al. 2005, Manoli et al. 2005, Kimura et al. 2005, Yu et al. 2010, Cachero et al. 2010, von Philipsborn et al. 2014). However, expression of neuronal transmitter types, including GABA, in the whole FruM positive cell population, as well as their contribution to circuit function has not been studied so far. GABAergic transmission is widespread in the *Drosophila* nervous system, with the GABA producing enzyme glutamic acid decarboxylase (Gad1), the ionotropic GABA receptor Rdl and the vesicular GABA transporter (vGAT) distributed in all major neuropil areas of the brain and ventral nerve cord (Jackson et al. 1990, Ffrench-Constant et al. 1991, Chen et al. 1994, Harrison et al. 1996, Enell et al. 2007). Single-cell transcriptomics analysis suggests that around 25% of *Drosophila* central brain neurons (Croset et al. 2018) and around 10% of the entire brain including optic lobes (Davie et al. 2018) are GABAergic. In the central nerve cord, around 38% of all neurons express Gad1, the marker enzyme for GABAergic neurons (Allen et al. 2020). In these datasets, visualized by SCope tool, co-expression of *Gad1* and *fruitless* is obvious in numerous neurons (Davie et al. 2018).

Previously, few GABAergic *fruitless* expressing (*fru+*) neurons have been investigated in detail in their role for courtship behavior. The GABAergic brain neuronal class mAL, for example, regulates the processing of gustatory information that informs the courting male of the sex of another fly. mAL provides inhibitory input to central courtship promoting P1 neurons. mAL is activated by leg gustatory neurons that sense male pheromones with courtship suppressing effect. It thus functions to prevent male-male courtship. Interestingly, mAL is also activated by different subsets of leg gustatory neurons that sense courtship-promoting female pheromones. Since the sensory neurons responsive to female pheromones activate both GABAergic mAL and the interneuron vAB3 that excites P1, its second function is probably to exert gain control in the response of P1 to courtship stimulating female pheromones (Kallmann et al. 2015, Clowney et al. 2015).

A more comprehensive view of how GABAergic inhibition shapes the entire behavioral sequence of male courtship, however, is missing. Moreover, it has never been addressed how GABAergic neurons contribute to generating the wing song motor pattern, a central element of male courtship whose correct execution strongly affects copulation success.

Here, we used RNAi mediated knockdown of Gad1 or Rdl expression in *fru+* neurons and asked how GABAergic signaling in the male specific circuit affected courtship. We further provide an anatomical description of all *Gad1+* and *fru+* neurons in brain and ventral nerve cord. We test how brain versus ventral nerve cord populations contribute to the various behavioral phenotypes overserved after disruption of GABAergic signaling. By focusing on courtship song patterning, we analyze with recording and playback experiment the causal relationship between disrupted song patterns and copulation success.

We find that the depletion of Gad1 and Rdl in *fru+* neurons leads to very similar specific defects in hierarchical organization and motor coordination of male behavior, preventing the animals from achieving copulation. Copulation is frequently attempted, but not in the appropriate proximity and orientation to the female. Males sing with both wings. Upon closer inspection of the acoustic signals generated by wing vibration, we find that knockdown of Gad1 or Rdl leads to severe alterations of the courtship song pattern. The sine song mode is strongly reduced and the pulse song mode is altered in pulse structure and spacing. This strongly altered song fails to stimulate the receptivity of wild type females. Wild-type song, however, cannot rescue the copulation defect of knockdown males, indicating that additionally to song, defects in copulation behavior prevents them from mating. Apart from the defects in the execution of the different courtship steps, Gad1 and Rdl depleted males are unable to reduce courtship after negative conditioning with unreceptive females. All observed singing and song patterning phenotypes result from depletion of GABAergic signaling in the ventral nerve cord. In contrast, manipulation of GABAergic neurons in the brain affects courtship conditioning.

Our study illustrates that GABAergic *fru+* neurons that inhibit *Rdl+ fru+* neurons tune many aspects of courtship behavior. We locate the circuit motifs for unilateral wing usage, sine song, pulse length and pulse spacing, as well as coordination of intromission in the ventral nerve cord. In the future, it will be interesting to identify the cellular identity and the connectivity of these motifs.

## Results

### GABAergic signaling in *fruitless* neurons is required for motor coordination of male courtship behavior

To assess the role of GABAergic signaling in male courtship behavior, we focused on the approx. 2% of adult male neurons expressing the sex determination factor FruM and analyzed males that had either the GABA producing enzyme Gad1 or the ionotropic GABA receptor Rdl depleted in all neurons expressing the *fruitless-GAL4* (*fru-GAL4*, Stocking et al. 2005) driver (*fru+* neurons) by RNAi mediated knockdown. In contrast to controls, knockdown males did not achieve copulation when paired with wild type virgin females for 20 min (**Fig. 1A**). This was not due to low courtship intensity, since knockdown male attempted copulation more often than controls (**Fig. 1B**). When we scrutinized the observed copulation attempts, however, we saw that knockdown males had a lower proportion of appropriate copulation attempts, i.e. attempts that were directed at the female abdomen and performed in the close vicinity of the female (**Fig. 1B, C**). Next, we asked if another central courtship element, male wing song, was affected by depletion of GABAergic signaling. Knockdown males extended their wings as often or, in the case of Rdl knockdown, more often than control males (**Fig. 1E**). The majority of the knockdown male wing extensions differed markedly from controls in such that the male extended two wings instead of one (**Fig. 1F, G**). Next, we wondered if knockdown males would abstain from singing when control males naturally do so, namely after courtship conditioning. Control males sing only very little or not at all after being trained with a mated, unreceptive female for 1 hr. Males depleted of Rdl sang the same amount of pulse song with or without training. Males depleted of Gad1 showed a slight reduction of pulse song. Their learning index, however, was significantly lower than learning indices of control flies (**Fig. 1 H**).

**Figure 1.**
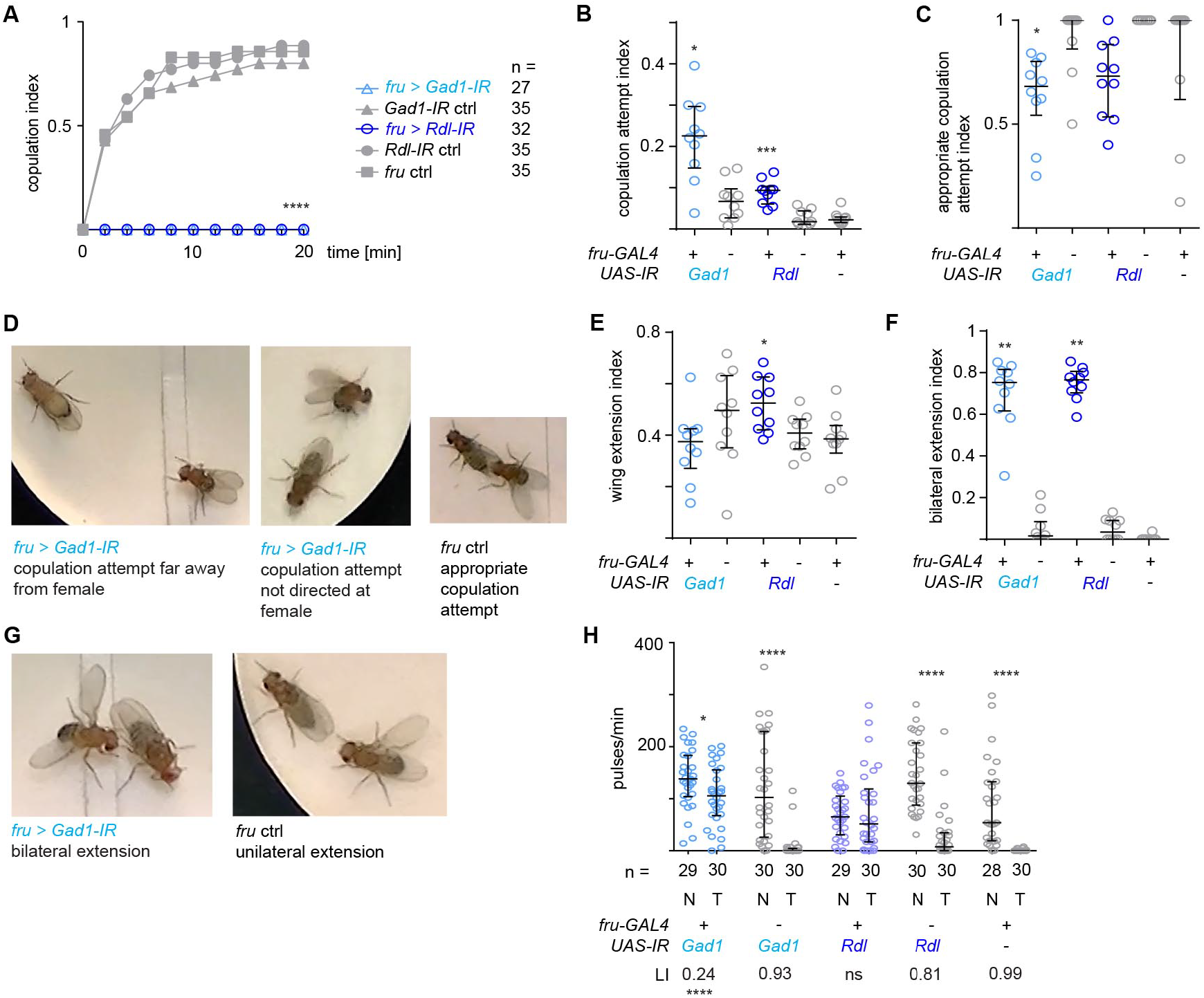
Depletion of GABAergic inhibition affects motor coordination of male courtship behavior. **A** Copulation index of male flies upon RNAi-mediated knockdown of *Gad1* or *Rdl* in *fruitless* neurons and respective genetic controls. ****p< 0.0001, Fisher exact text. n indicates number of flies tested. **B** Copulation attempt index (fraction of video frames in which the male attempted copulation) for knockdown and control males. **C** Appropriate copulation attempt index (fraction of copulation attempts which were directed at and in reach of the female) for knockdown and control males. **D** Examples of inappropriate copulation attempts of a *Gad1* depleted male and an appropriate copulation attempt of control male. **E** Wing extension index (fraction of video frames in which the male extended a wing) for knockdown and control males. **F** Bilateral wing extension index (fraction of wing extensions which were bilateral) for knockdown and control males. **G** Example of bilateral wing extension of a *Gad1* depleted male and a unilateral extension of control male. **H** The effect of courtship conditioning in knockdown and control males. For each genotype the amount of song (in pulses/min) toward a mated wild type female is shown for naïve (N) males and males trained (T) with a unreceptive mated wild type female. * p = 0.02, ****p<0.0001, Mann-Whitney test. The learning index of each genotype (LI) is given under the graph in the cases were significant learning occurred. ****p < 0.0001 indicates significant difference between LI of *fru > Gad1* and the LI of controls, permutation test. n indicates number of flies tested. **A-G:** All experiments are performed with wild type virgin females. **B, C, E, F**: Each knockdown genotype is compared to its corresponding *UAS-IR* control and the *fru* ctrl, *p<0.05, **p<0.005, Kruskal-Wallis test with Dunn’s multiple comparison, n = 10 males per genotype. In all scatterplots, each data point represent one fly, error bars indicate median with interquartile range.

We conclude that males depleted of GABAergic inhibition in *fru+* neurons do not copulate within normal time spans, fail to suppress inappropriate courtship attempts, extend two wings during singing and show no or strongly diminished reduction of singing after courtship conditioning.

### Patterning of courtship song depends on GABAergic signaling

The postural defect during singing led us to the question if depletion of GABAergic signaling in *fru+* neurons also affected the patterning of the acoustic properties of courtship song. Knockdown males reliably sang pulse song, albeit a lower amount than control males (**Fig. 2A**). As immediately apparent from sample oscillograms, song of males depleted for either Gad1 or Rdl in *fru+* neurons had marked and similar changes compared to song of control males (**Fig. 2B**). Sine song was strongly reduced (**Fig. 2C**) and pulse song was polycyclic, i.e. individual pulses increased in length and the number of underlying wing strokes (cycles) (**Fig. 2C**). While control pulse song is characterized by a high percentage of pulses which are spaced at 30-35 ms, the distribution of inter pulse intervals (ipis) of knockdown flies was less defined and broader, with less ipis in the 30-40 ms range (**Fig. 2E, F**). After knockdown of Gad1, but not Rdl, the carrier frequency of pulse and sine song increased (**Fig. 2G, H**). To confirm that these phenotypes resulted specifically from depletion of GABAergic signaling, we repeated knockdown experiments with independent RNAi lines against *Gad1* and *Rdl*, as well as an RNAi line against the vesicular GABA transporter *vGAT*. We reasoned that knockdown of the latter, which is essential for packaging GABA into synaptic vesicles, would lead to similar phenotypes than interference with GABA synthesis. Males depleted for GABAergic signaling with these alternative RNAi lines showed very similar song defects to the ones described above (**Supplementary Fig. 1**). We conclude that GABAergic, *fru+* neurons and *fru+* neurons expressing Rdl are required for correct patterning of courtship song.

**Figure 2.**
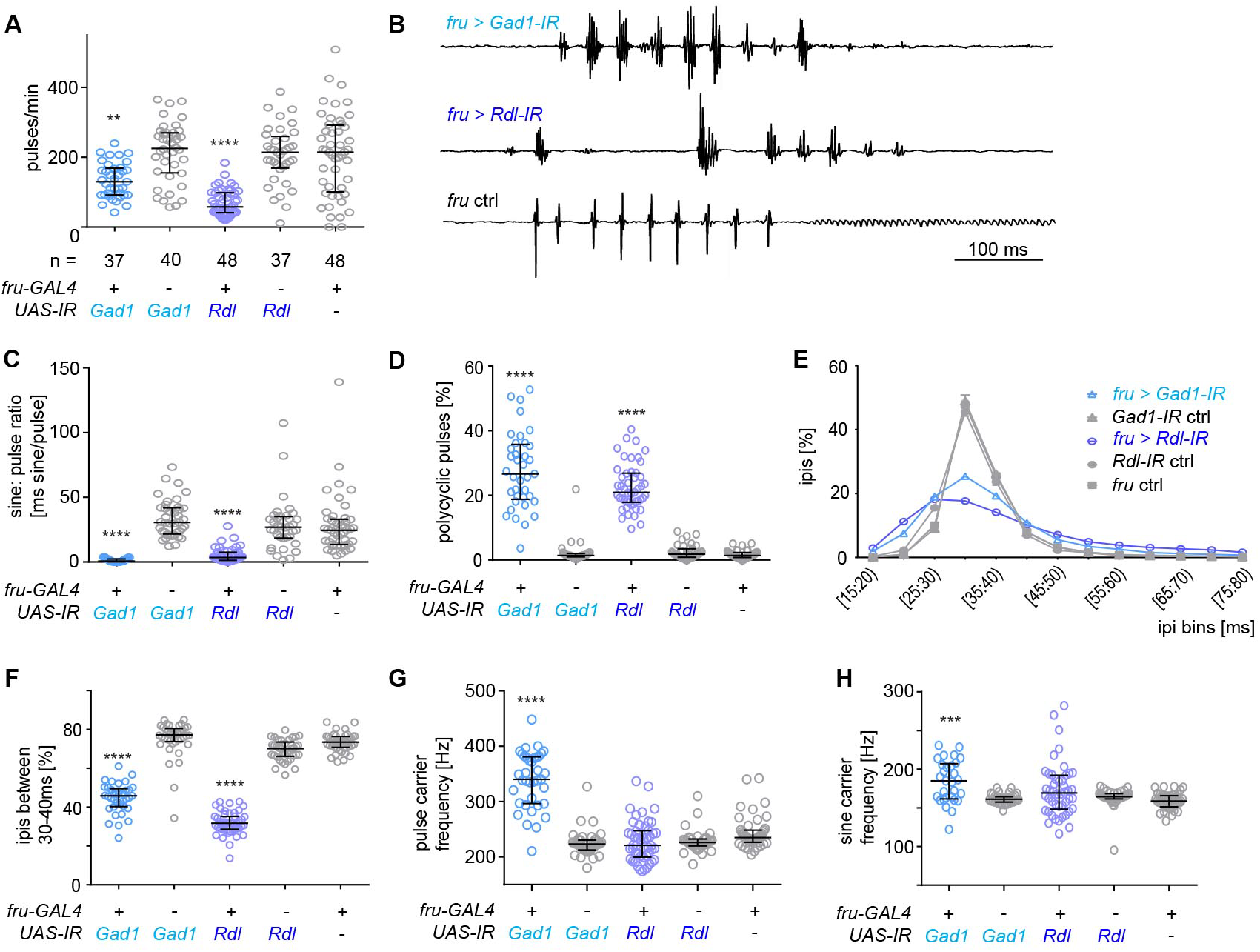
GABAergic signaling in *fruitless* neurons is required for courtship song patterning. **A** Amount of courtship song (in pulses/min) displayed toward a virgin wild type female of male flies upon RNAi-mediated knockdown of *Gad1* or *Rdl* in *fruitless* neurons and respective genetic controls. n indicates number of flies tested. **B** Examples of oscillogramms of courtship song from knockdown males and a control male. **C** Amount of sine song (sine: pulse ratio) for knockdown and control males. **D** Amount of polycyclic pulses (% of pulses with more than 2 cycles) in the song of knockdown and control males. **E** Distribution of inter pulse intervals (ipis) in the song of knockdown and control males. For each genotype, the mean percentage of ipis in the corresponding bin among flies and the standard error of the mean is plotted. **F** Percentage of inter pulse intervals (ipis) within the 30 – 40 ms range in the song of knockdown and control males. **G** Median pulse song carrier frequency for knockdown and control males. **H** Median sine song carrier frequency for knockdown and control males. **A, C, D, F-H**: Each knockdown genotype is compared to its corresponding *UAS-IR* control and the *fru* ctrl, *p = 0.017,***p = 0.0007, ****p< 0.0001, Kruskal-Wallis test with Dunn’s multiple comparison; n in **C-H** is the same as in **A**. All experiments are performed with wild type virgin females. In all scatterplots, each data point represent one fly, error bars indicate median with interquartile range.

### Distribution of *fru+* GABAergic neurons

Having established the functional relevance of *fru+* GABAergic neurons for behavioral coordination and song patterning during male courtship, we next were interested to identify specific neurons responsible for the observed defects. Since the neurotransmitter identity of *fru+* neurons has not been comprehensively mapped, we first identified all *fru+* GABAergic neurons marked by coexpression of *fru-GAL4* and *Gad1-Lex* (Diao et al. 2015) transgenes (**Fig. 3A**). In the brain, such cells were present in seven distinct clusters, some of which have been previously named and described (**Fig. 3B, C**). In the ventral nerve cord (VNC), *fru+ Gad1+* neurons were less clustered and more variable regarding the position of cell bodies, but could be grouped as belonging to four different regions (**Fig. 3B, D**). We counted the anatomically defined neuronal classes/groups, which comprised 1-35 cells each, and found 130 ± 19.8 (n = 5, s.d.) neurons in the brain and 89.6 ± 20.5 (n = 8, s.d.) neurons in the VNC, that is, 11 subpopulations of approx. 220 *fru+ Gad1+* neurons in the entire central nervous system. Functions of some *fru+* GABAergic brain neurons are known, but neurons in the VNC have not been explored before (**Table 1**).

**Figure 3.**
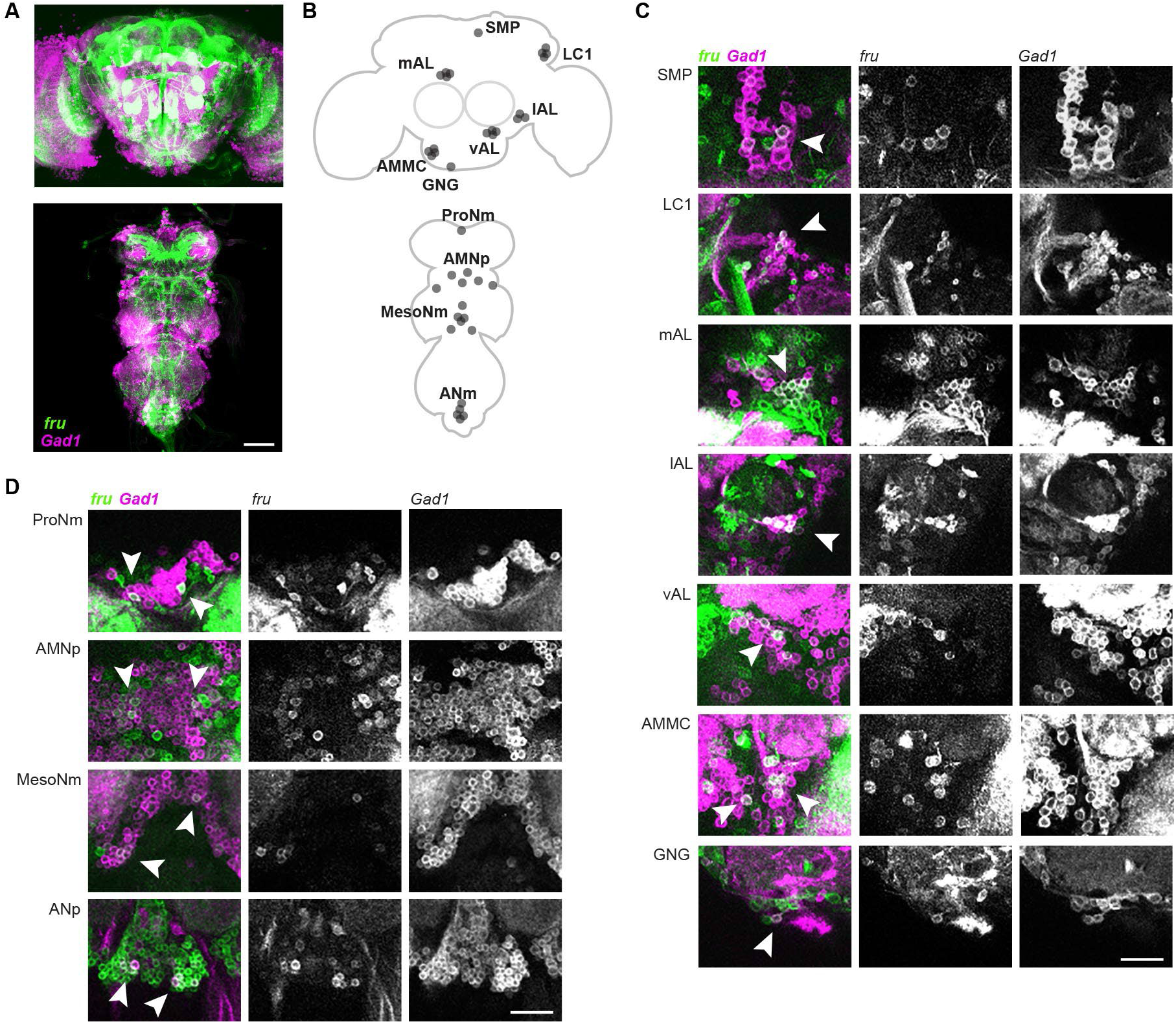
Distribution of *fru+* GABAergic neurons in brain and VNC. **A** *fru+* GABAergic neurons identified by coexpression of *fru-GAL4* (green, labeled with Gfp) and *Gad1-Lex* transgenes (magenta, labeled with Tomato) in brain and VNC. **B** Schematic of *fru+* GABAergic neurons. **C** *fru+* GABAergic neuronal classes in the brain. **D** *fru+* GABAergic neuronal classes in the ventral nerve cord. In **C** and **D**, arrowhead point to coexpression. Scale bars: 100 μm. SMP, superior medial protocerebrum; LC1, LC1 neurons; mAL, mAL neurons; IAL, lateral antennal lobe; vAL, ventral antennal lobe; AMMC, antennal mechanosensory and motor center; GNG, gnathal ganglia; ProNm, prothoracic neuromere; ANMp, accessory mesothoracic neuropil; MesoNm, mesothoracic neuromere; ANm, abdominal neuromeres. Brain region nomenclature in accordance with (Ito et al. 2014), VNC region nomenclature in accordance with (Court et al. 2017). See also Table 1 for cell counts and comments.

**Table 1.**
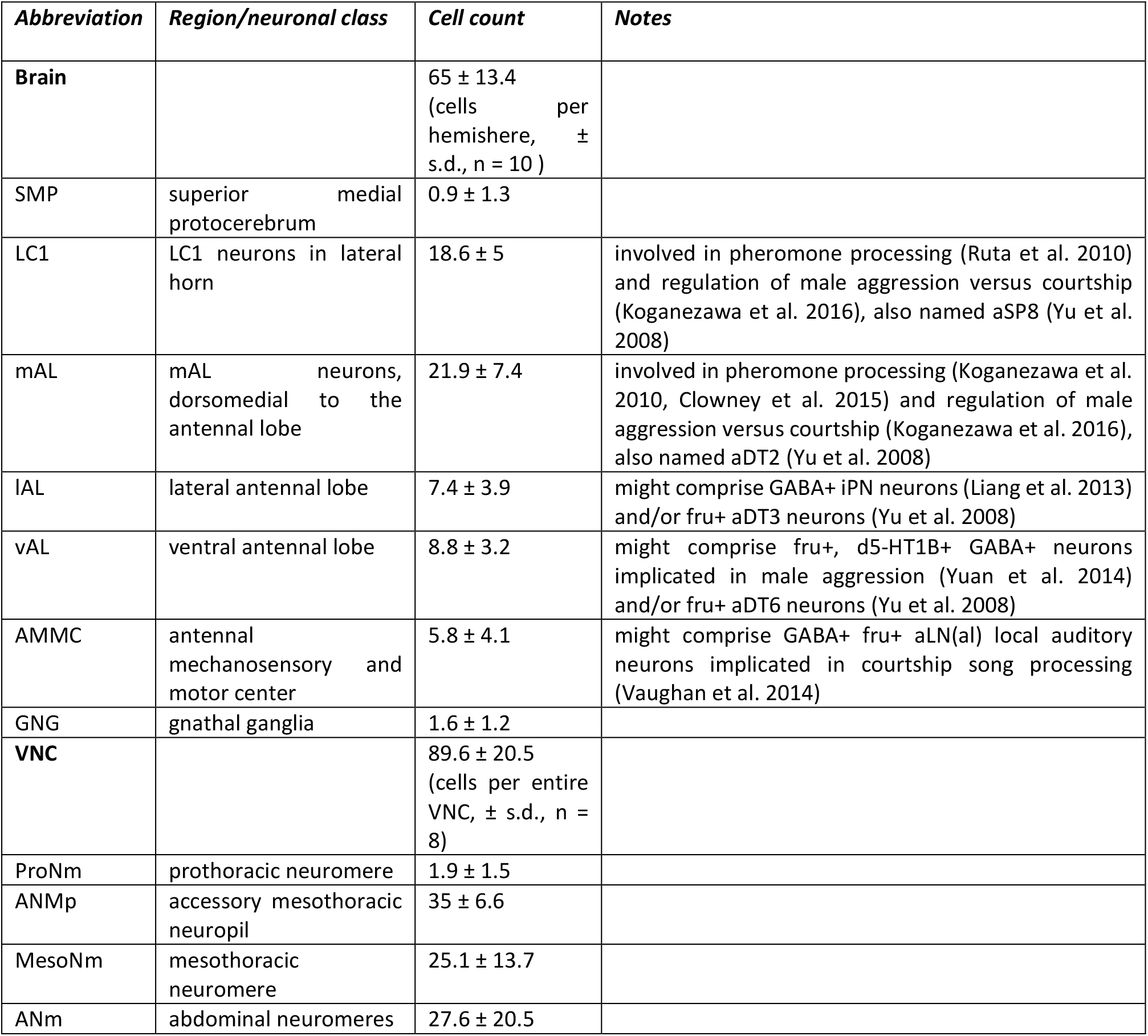
*Gad1+fru+* neurons in the central nervous system. Cells coexpressing *fru-GAL4* and *Gad1-LexA* in different brain and VNC regions of male flies. Brain region nomenclature in accordance with (Ito et al. 2014), VNC region nomenclature in accordance with (Court et al. 2017). Brain neurons were observed in more discrete clusters and counted by hemispheres, VNC neurons had more variable and dispersed positions and were counted per entire tissue. 130 ± 19.8 neurons were counted in the entire brain (n = 5 brains, s.d.). For cell locations and exemplary micrographs, see Figure 3.

### *fru+* GABAergic neurons in the VNC are required for song patterning

To distinguish between the functional roles of *fru+* GABAergic neurons in the brain versus in the VNC, we restricted the knockdown of Gad1 and Rdl to each of the two tissues using the brain specific *otd-FLP* transgene in combination with different FLP recognizable Gal80 cassettes. As visualized by Gfp expression, this approach resulted in the respective expression domains, allowing us to probe *fru+* neurons in the brain separately from *fru+* neurons in the VNC (**Fig. 4A**). When either Gad1 or Rdl was knocked down in the VNC only, flies displayed bilateral wing extension during singing. In contrast, brain restricted knockdown did not lead to any bilaterality phenotype. Likewise, the alterations in acoustic song properties seen upon knockdown of Gad1 or Rdl in *fru+* neurons of the entire nervous system were mostly replicated by VNC restricted, but not brain restricted knockdown (**Fig. 4B**). Gad1 and Rdl were required in *fru+* VNC, but not brain neurons for short-term copulation success (**Fig. 4C**). Given the changes in song structure upon manipulation of *fru+* VNC neurons, we wondered if rejection of altered song by females was causing the observed defects in copulation success. To test this hypothesis, we experimentally separated song phenotype and male genotype by amputating both wings of wild type male flies and playing back song from Gad1 knockdown males or wild type male song while the amputated males courted virgin females. Wild type song, but not structurally changed song from *fru>Gad1-IR* knockdown males could rescue the short-term copulation success of wing-amputated males. Song from knockdown males was equally inefficient as no playback at all, with both conditions leading to no copulations among 40 couples after 30 min (**Fig. 4D**). This indicates that female receptivity is not sufficiently stimulated by courtship song altered by Gad1 depletion. When wild type song was played back during courtship of wing-amputated males with Gad1 or Rdl knockdown in *fru+* VNC neurons, these males remained unsuccessful in obtaining copulations (**Fig. 4E**). We conclude that defect song upon Gad1 or Rdl knockdown in *fru+* VNC neurons is sufficient to cause a defect in copulation success. However, *fru+* GABAergic neurons in the VNC are also required for copulation in some other way, playing an additional, song-independent role.

**Figure 4.**
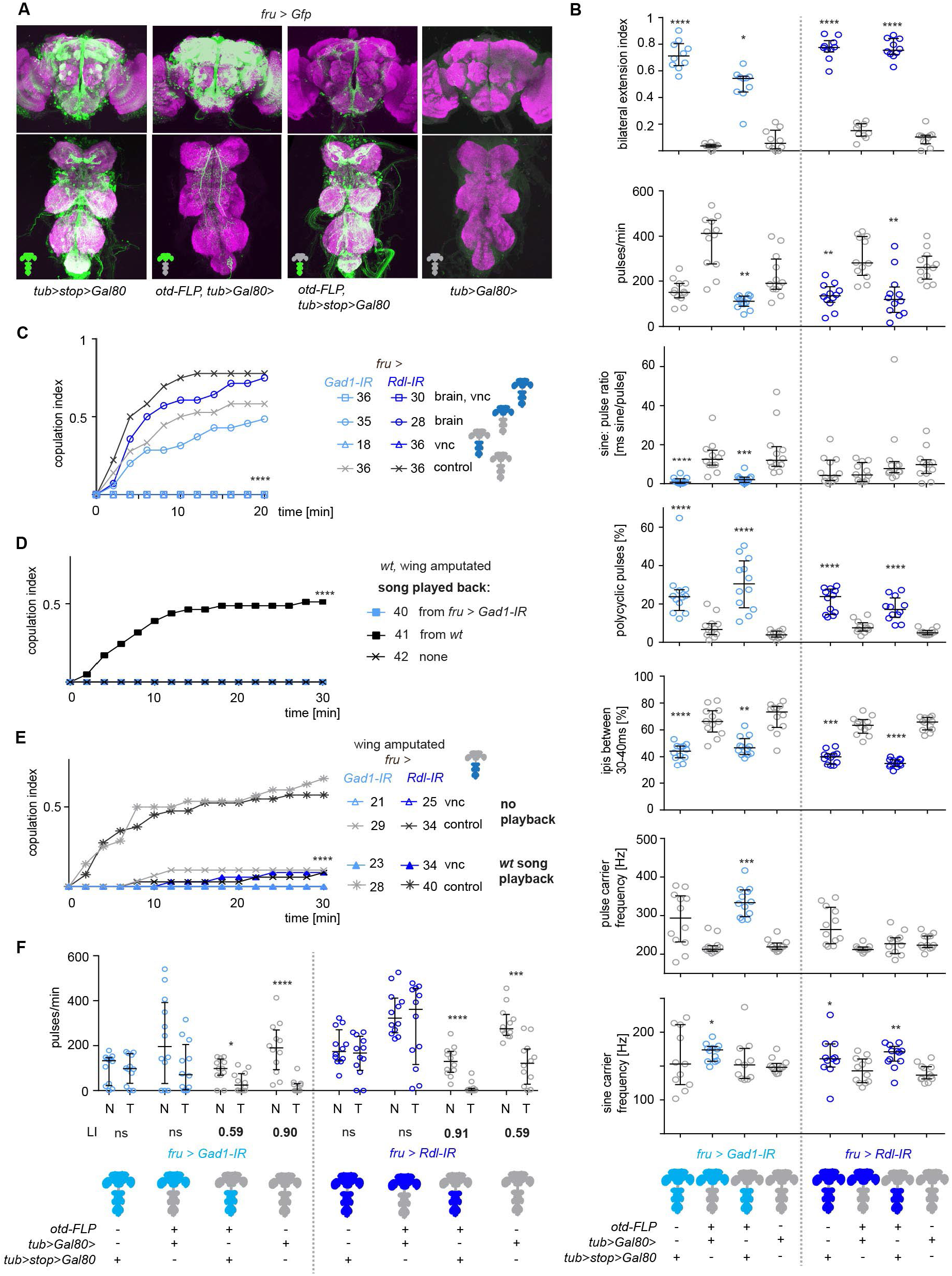
Distinct roles of *fru+* GABAergic neurons in brain and VNC. **A** Representative brain or VNC-restricted transgene expression in *fru+* GABAergic neurons by combining brain specific *otd-FLP* with different constructs of the GAL4 repressor GAL80 and genetic controls. Neuronal arbours in the VNC of the brain-specific *otd-FLP, tub>stop>GAL80* specimen are from descending neurons, neuronal arbours in the brain of the VNC-specific *otd-FLP, tub> GAL80>* specimen are from ascending neurons and neurons in the gnathal ganglia.GFP expression in green, neuropil anti-bruchpilot staining in magenta, scale bar: 100 μm. **B** Effects on song wing usage and song patterning of brain or VNC-specific RNAi-mediated knockdown of *Gad1* or *Rdl* in *fruitless* neurons and respective genetic controls. n =10 per genotype for bilateral extension index, n = 12 flies per genotype for song parameters. Full and partial knockdown genotypes are compared to the fully repressed *tub>gal80>* controls, *p < 0.05, **p < 0.005, ***p < 0.0005, ****p< 0.0001, Kruskal-Wallis test with Dunn’s multiple comparison. **C** Copulation index of male flies upon RNAi-mediated knockdown of *Gad1* or *Rdl* in *fruitless* neurons and respective genetic controls. ****p< 0.0001, Fisher exact test. **D** Copulation index of wing-amputated *wt* male flies supplemented with played back courtship song from a *fru > Gad1-IR* or *wt* male. ****p< 0.0001, Fisher exact test. **E** Copulation index of wing-amputated males with VNC-specific knockdown and controls with and without played back courtship song from *wt* male. ****p< 0.0001, Fisher exact test. **F** Effect of courtship conditioning in knockdown and control males. For each genotype the amount of song (in pulses/min) toward a mated wild type female is shown for naïve (N) males and males trained (T) with a unreceptive mated wild type female. * p = 0.02, ****p<0.0001, Mann-Whitney test. The learning index of each genotype (LI) is given under the graph in the cases where significant learning occurred. n = 12-15 flies per group/genotype. **B-E**: All experiments are performed with wild type virgin females. **C-E**: n, number of flies tested, is indicated in the legend. **B-F**: Blue colored parts in the nervous system cartoons indicate expression of the *IR* transgenes mediating knockdown. In B and F, genotypes with significant difference to the respective *tub>Gal80>* control (expression of *IR* transgene repressed in the whole nervous system) are indicated by blue colored data points.

Lastly, suppression of singing after courtship conditioning depended on Gad1 and Rdl expression in *fru+* brain neurons and was not affected by Gad1 or Rdl depletion in *fru+* VNC neurons (**Fig. 4F**). In summary, different courtship and mating defects upon depletion of GABAergic signaling in fru+ neurons, are due to different groups of neurons. Main song patterning defects (bilateral wing usage, reduction of sine song, polycyclicity and broadening of the ipi range) and short-term copulation failure result from impairing GABAergic *fru+* neurons in the VNC. Defects in courtship conditioning, i.e. loss of the reduction of song after training, result from impairing GABAergic *fru+* neurons in the brain.

## Discussion

*Drosophila* male courtship is an excellent model system of how complex behavior is orchestrated by a male specific set of interconnected neurons, the *fru+* circuit (Yu et al. 2008). So far, publicly available connectomes of the *Drosophila* nervous system from electron microscopy (EM) reconstructions come from female animals (Zheng et al. 2018, Phelps et al. 2018, Scheffer et al. 2020), but rapid advance in this area will soon allow for in more detail the synaptic connectivity of identified *fru+* neurons. Computational tools allow for prediction of neurotransmitter type from EM images (Eckstein et al. 2020), the analysis of large light microscopy expression datasets (Panser et al. 2016) and the cross-comparison of EM and light microscopy data (Costa et al. 2016). In the face of this wealth of anatomical data, it is critical to mechanistically link the architecture of circuit motifs to behavior. Based on our analysis of depletion of GABAergic signaling the *fru+* circuits in brain and ventral nerve cord, we propose several hypotheses of how inhibitory motifs shape critical aspects of male courtship behavior and thus are required for mating success.

### Inhibitory signaling is required for male copulation success

We find that RNAi mediated knockdown of Gad1 or Rdl in *fru+* neurons of the ventral nerve cord prevents copulation of the manipulated male with a wild type female. Absence of copulation could stem from either the lack of female receptivity or the lack of male courtship intensity, motor skill and coordination. We find indication that both is the case. The lack of inhibition alters the species-specific pattern of the courtship song to such an extent that playback of the acoustic signal, in contrast to control song, cannot stimulate receptivity of wild type females (**Fig. 4D**). If defect song was the main cause of preventing knockdown males from copulating, playback of *wt* control song to wing amputated animals should rescue the phenotype. We only observe such rescue in control males with functional GABAergic signaling (**Fig. 4D**). Judged from the frequency of wing extensions and the number of copulation attempts (**Fig. 1B, E**), knockdown of *Gad1* or *Rdl* does not decrease overall courtship intensity, but slightly increases it. The males are clearly capable of bending their abdomen in the typical copulation attempt posture. However, they attempt copulation when it is futile, i.e. when the female is far away or at an inappropriate angle (**Fig. 1C, D**). This suggests that in normal courtship, copulation attempts are tightly gated by the spatial relationship of the two sexes. We hypothesize that disinhibition of copulation attempt commend neurons, e.g. aSP22 (McKellar et al. 2019) could allow the male to perform precisely timed attempts. A GABAergic interneuron receiving proximity information might inhibit a GABAergic gating neuron that is connected to ventral nerve cord arbors of aSP22 (**Fig. 5A**). The proximity and positional information might be derived from both visual and olfactory stimuli. These modalities have previously been shown to act redundantly to guide the defined position of the courting male behind the female (Kimura et al. 2015).

**Figure 5.**
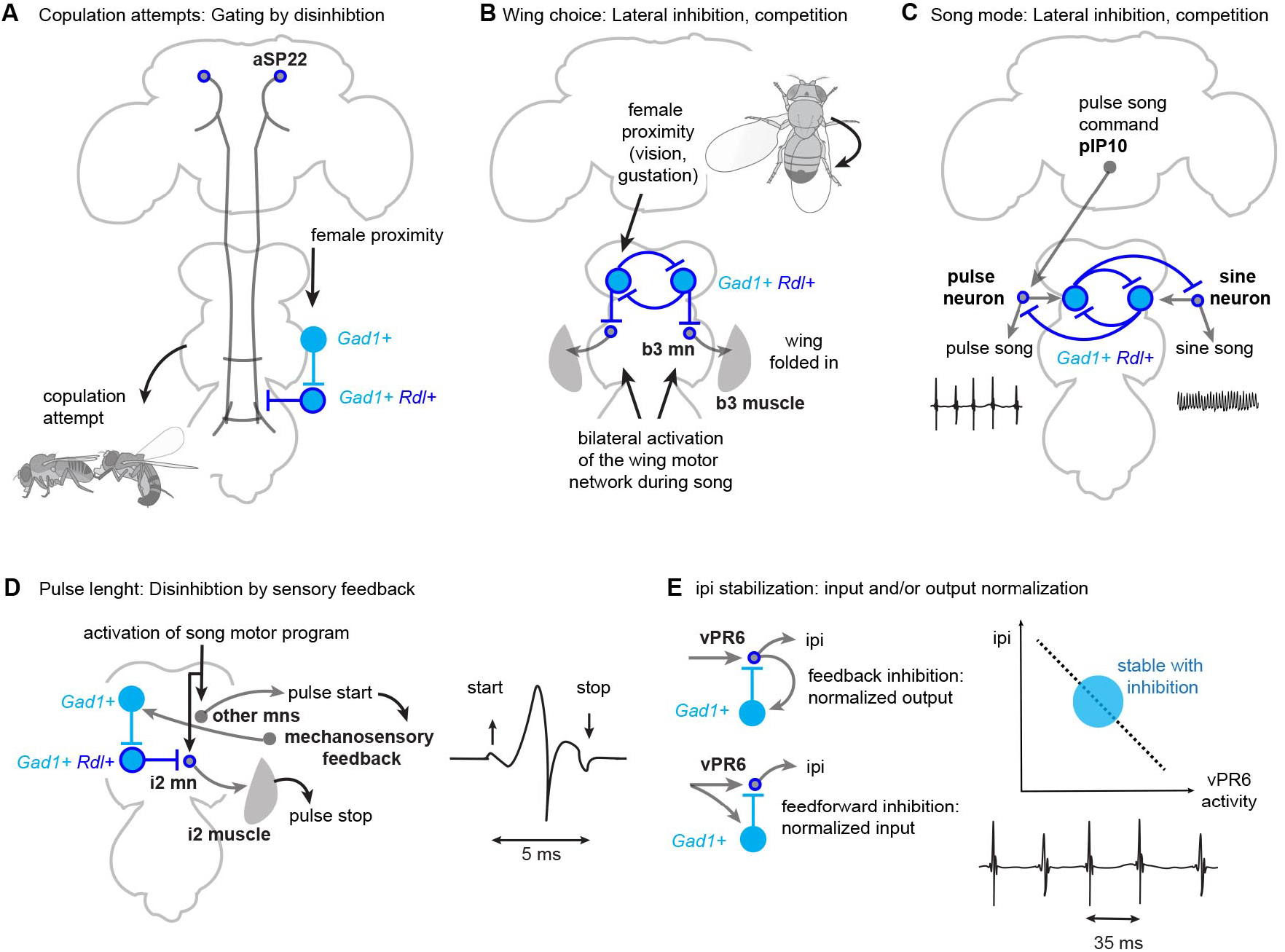
Proposed inhibitory circuit motifs for aspects of courtship behavior. **A** Fast and precisely timed copulation attempts could arise by disinhibition of the command neuron for copulation attempts, aSP22 (McKellar et al. 2019). Loss of inhibition leads to inappropriate copulation attempts. **B** Circuit model for wing choice during song and its dependence on a circuit motif with reciprocal inhibition. Lateralized information about female proximity inhibits the ipsilateral b3 mn and disinhibits the contralateral b3 mn. Contralateral b3 mn and b3 muscle activation draws in the contralateral wing. The lateral inhibition motif allows for stable wing choice in the absence of strongly lateralized sensory input. Loss of inhibition elads to bilateral wing usage during singing. **C** Model proposed by Roemschied et al. 2021 for alternation of pulse and sine song, adapted to a version based on *Gad1+ Rdl+ fru+* neurons. Both song modes are elicited by the pulse command neuron pIP10. Sine song neurons are active by rebound excitation after feedback inhibition has terminated a pulse song train. Loss of inhibition leads to loss of sine song. **D** Restriction of pulse length by mechanosensory feedback from pulse start, activating a disinhibition motif gating i2 mn activity. i2 mn and i2 muscle activity stop wing movement (O’Sullivan et al. 2018). Loss of inhibition leads to long, polycyclic pulses. **E** Restriction of ipi variability by normalization of vPR6 activity. vPR6 activity is inversely correlated with ipi (von Philipsborn et al. 2011). Stabilization of vPR6 activity could arise from simple feedback or feedforward inhibition. Loss of inhibition leads to a a broadening of the IPI distribution. Expression of Gad1 is indicated in Cyan, expression of Rdl in Blue. aSP22, pIP10, vPR6 and the yet unidentified *Gad1+* and *Gad1+ Rdl+* cells depicted are *fru+*.

### GABAergic inhibition in the brain mediates courtship conditioning

GABAergic inhibition in *fru+* circuit motifs in the brain has previously been suggested to act as gain control, normalizing the response to excitatory courtship stimuli (Kallmann et al. 2015, Clowney et al. 2015). One might therefore predict excessive courtship upon the loss of inhibition. Interestingly, we do not see a large increase of courtship singing upon depletion of GABAergic signaling (**Fig. 1E, 2A**). Loss of inhibition however lead to a strong increase in the amount of song in a situation where control males sing little, i.e. after aversive conditioning with an unreceptive female. Courtship conditioning is often used as a paradigm for studying learning and memory (Griffith and Ejima 2009, Raun et al. 2021). The inability of knockdown males to undergo courtship conditioning in our experiments can be interpreted as various, not mutually exclusive defects: the inability to perceive unsuccessful courtship as aversive, the inability to form a memory of the aversive experience and the inability to recruit the appropriate motor program (courtship suppression) upon retrieval of the memory. Since little is known about GABAergic signaling in the circuits for courtship conditioning, we here remain agnostic about the underlying causes of this phenotype.

### GABAergic control of unilateral wing usage

Normal male courtship song is accompanied by unilateral wing extension. Depletion of Gad1 or Rdl leads to the prevalence of bilateral wing extension. Wing choice during courtship singing depends on both gustatory and visual input (Koganezawa et al. 2010, Ribeiro et al. 2018). During unilateral singing, the wing control muscle b3 is active on the side of the folded wing, suggesting that its innervating motor neuron b3 mn prevents extension and full vibration of the wing not used in singing (O’Sullivan et al. 2018, Swain and von Philipsborn 2021). We hypothesize that reciprocal inhibition of *Rdl+ Gad+ fru+* interneurons that also inhibit b3 mn ensure unilateral wing usage (**Fig. 5B**). These neurons are predicted to receive input from the above-mentioned sensory pathways.

A similar network motif with reciprocal inhibition has been shown to direct the lateralized escape response of zebrafish larva (Koyama et al. 2016) and might be a widespread architecture allowing behavioral choice during asymmetric motor behavior of bilateral animals.

### Courtship song patterning by GABAergic inhibition

Rhythmically structured motor programs support many essential behaviors such as locomotion, breathing and eating (Kiehn and Kullander 2004, Sakurai and Katz 2015, Paton and Buonomano 2018, Mantziaris et al. 2020). They are also important in animal communication, conferring relevant information such as species identity and sender fitness to signals exchanged between conspecifics (Greenfield 2002). *Drosophila* male song has an organized, precisely timed structure and serves as good model for studying circuit mechanisms underlying pattern generation (Swain and von Philipsborn 2021). A small set of wing motor neurons has been shown to control main song parameters: the prevalence of the two song modes (sine song and pulses song), the length of pulses, and the spacing of the pulses (ipi), (O’Sullivan et al. 2018). We find that inhibitory interneurons in the ventral nerve cord are required for sine song production, restriction of the pulse length to 1-2 wing beat cycles and the timing of the IPI (**Fig. 2A-F**, **Fig. 4B**). A few *fru+* interneurons in the ventral nerve cord that affect song structure have been identified (von Philipsborn et al. 2011, Shirangi et al. 2016), but to our knowledge none of these identified neurons has been shown to be GABAergic. A recent model by Roemschied et al. (2021) suggests that inhibitory neurons are at the core of pulse and sine songs sequencing. A pair of neurons that reciprocally inhibit each other as well as either a pulse or a sine activating neuronal element —could generate and alternation of the both song modes (**Fig. 5C**). Shiozaki et al. (2022) find that overlapping neuronal populations are active during sine and pulse song, with less neurons active during sine and additional neurons recruited during pulse. This could be interpreted as sine song relying on inhibition of a pulse specific interneuron population. In both models, loss of neuronal inhibition would lead to loss of sine, but not pulse song, as long as excitatory input from song promoting brain neurons is intact.

Additional to the severe reduction of sine song, we observe the appearance of pulse song with long, polycyclic pulses upon knockdown of Gad1 or Rdl (**Fig. 2B, D, 4B**). This phenotype resembles the one seen upon silencing of the motor neuron innervating the wing control muscle i2, i2 mn. The i2 muscles has been proposed to act as a “pulse stopper”, actively terminating the vibration of the wing after 1-2 cycles of up- and downward movement (O’Sullivan et al. 2018). Moreover, it has been suggested that such a pulse stop mechanisms could arise from mechanosensory feedback (Ewing 1979). Here, we propose that disinhibition of i2 mn by *Gad+ Rdl+ fru+* interneurons, activated by mechanosensory feedback from the initiation of the wing beat, prevents normal song pulses from escalating into a polycyclic pattern (**Fig. 5D**). Such an escalation is observed when the inhibitory motif is impaired, since wing power muscles are stretch activated and support longer wing oscillations once the thorax is deformed by a wing deflection (Dickinson and Tu 1997).

Wild type courtship pulses occur in trains where they are spaced at approximately 35 ms at 25°C. While in control song, around 80% of all inter pulse interval (ipis) are 30-40 ms long, depletion of Gad or Rdl leads to an ipi distribution that is more variable, but has a similar mode value (**Fig. 2E, E, 4B**). Very little is known about circuit mechanism that control ipi. Possibly, ipi timing could be disrupted if it relies on correctly timed firing of neurons involved in the length and cyclicity of a single pulse. Previous work has shown that increasing activation strength of the *fru+* interneuron vPR6 leads to decreased ipis of the elicited pulse trains (von Philipsborn et al. 2011). vPR6 does not express Gad1 (**Fig. 4A, B**). If vPR6 firing rates have to be within a certain range to maintain the normal ipi distribution, its output and/or excitatory input could be normalized and stabilized by simple feedback inhibition or feedforward inhibition circuit motifs (**Fig. 5E**). Future identification of genetic lines for the specific manipulation of individual *fru+* GABAergic neuronal classes among the around 90 ventral nerve cord cells (**Table 1**) will open the possibility to study song patterning in more detail.

Nervous systems generate behavioral output in accordance to the internal state of the animal and process external stimuli relevant to this state. Neuronal inhibition enables them to do so in an efficient and adaptive way. GABAergic inhibition fine-tunes and organizes male *Drosophila* courtship behavior on many levels. Upon depletion of inhibitory, males still perform all steps of the behavioral sequence in response to females. However, the motor execution is inaccurately timed and uncoordinated, severely reducing reproductive success. Further studies, helped by male nervous system connectomes, should identify the cellular identities of the *fru+, Gad+* and/or *Rdl+* interneurons underlying the here described behavioral phenotypes and test our proposed motif models for gating of copulation attempts, unilateral wing, sequencing of sine and pulse song and pulse timing.

## Material and Methods

### Transgenic *Drosophila* lines

*Drosophila* lines used and their origin are listed in **Table 2**. For control genotypes, GAL4 and UAS lines were crossed with wild type flies. For RNAi mediated knockdown with lines from the VDRC libraries, *UAS-Dcr2* was used to enhance knockdown efficiency (Supplementary Figure 1). Flies were grown on regular food (water, cornmeal, oatmeal, sucrose, yeast, agar, acetic acid, preservative methyl-4-hydroxybenzoate) at 25°C, 60% humidity, and a l2h/l2h light-dark cycle.

**Table 2.** Transgenic *Drosophila* lines.

| **Genotype** | **Short name** | **source** | **RRID** |
| --- | --- | --- | --- |
| *fruitless-GAL4* | *fru-GAL4* | Gift from Barry Dickson | BDSC_66696 |
| *y v; P{TRiP.HMC03350}attP40* | *UAS-IR Gad1, Gad1-IR* | Bloomington Drosophila Stock Center (BDSC) | BDSC_51794 |
| *w<sup>1118</sup>; P{GD8508}v32344* | *UAS-IR Gad1\** | Vienna Drosophila Resource Center (VDRC) | SCR_013805 |
| *y sc v sev; P{TRiP.HMC03643}attP40* | *UAS-IR Rdl, Rdl-IR* | BDSC | BDSC_52903 |
| *w<sup>1118</sup>; P{GD4609}v41103* | *UAS-IR Rdl\** | VDRC | SCR_013805 |
| *P{KK101500}VIE-260B* | *UAS-IR vGAT* | VDRC | SCR_013805 |
| Wild type Canton S | *wt* | Gift from Barry Dickson | n.a. |
| *w<sup>1118</sup>; P{UAS-dicer2, w[+]}* | *UAS-Dcr2* | VDRC | SCR_013805 |
| *w-; UAS-mCD8-Gfp; +* | *UAS-Gfp* | Gift from Barry Dickson | n.a. |
| *w-; LexAop-mCD8-tomato; +* | *LexAop-tomato* | Gift from Barry Dickson | n.a. |
| *w; Mi{Trojan-lexA:QFAD.2}Gad1[MI09277-TlexA:QFAD.2]* | *Gad1-LexA* | Gift from Ben White | BDSC_60324 |
| *w; Otd-nlsFLPo.attP40; +* | *Otd-FLP* | Gift from David Anderson | n.a. |
| *w; +; tubP-FRT-stop-FRT-GAL80* | *tub>stop>Gal80* | BDSC | BDSC_39213 |
| *w; +; tubP-FRT-GAL80-FRT* | *tub> Gal80>* | Gift from Barry Dickson | n.a. |

### Behavioral assays and data evaluation

For all behavioral assays, if not otherwise indicated, males were individually aged for 4-7 days and paired with 4-7 d old CS virgins that were aged in groups of 10-20 flies.

For evaluating copulation success, pairs were videotaped for 20 min in chambers of 1 cm diameter. For evaluating wing extension and copulation attempts, pairs were videotaped in chamber of 1.7 cm diameter with beveled walls, which prevented flies from walking at the side walls of the chambers and allowed for continuous top view. Wing extensions were scored for the first 5 min, every 2.5 s, i.e. in approximately 120 frames per pair. The wing extension index was calculated as the fraction of frames in which the male extended at least one wing at an angle of more than 30°. The bilateral extension index was calculated as the fraction of wing extensions during which the male extended both wings at an angle of more than 30°. The large angle extension index was calculated as the fraction of wing extensions during which the male extended at least one wing at an angle of more than 85°. Copulation attempts were scored for the first 2 min, every 0.5 s, i.e. in approximately 120 frames per pair. The copulation attempt index is the fraction of frames in which the male attempted copulation, characterized by pronounced abdominal bending. The appropriate copulation attempt index is the fraction of copulation attempts during which the male pointed his abdomen toward the female genitalia and was not further than 1 fly length away from the female. For courtship suppression assays, males were paired for 1 hr with a mated CS female in a chamber of 1 cm diameter (trained group) or left alone for 1 hr in the chamber (naïve group). Males were then transferred into song recording chambers with a new mated female and courtship song was recorded for 4 min. Learning and suppression of song was judged to have occurred when the amount of pulse song was significantly different in trained and naïve groups. The learning index (LI) for each genotype was calculated as the relative reduction of pulse song in trained males as compared to naïve males: (mean (pulses/min)^Naive^-mean (pulses/min)^Trained^)/ mean (pulses/min)^Naιve^. To test for differences between the LIs of two genotypes, pulse/min values from individual flies from trained groups were permuted between genotypes, and the same was done for naïve groups (1000000 permutations). The p value gives the probability for obtaining a difference in Lis equal to or larger than the experimentally observed one.

Song recording and evaluation was performed as described previously (O’Sullivan et al. 2018). In brief, we used a multi-channel array designed by the Stern lab, detected pulse and sine song with a custom MATLAB script (Arthur et al. 2013) and used visualization of oscillograms to manually correct automated detection. Each datapoint corresponds to an approximately 4 min long recording of one male fly. The sine: pulse ratio was calculated by dividing the total amount of sine song in ms by the total number of pulses. Pulse cyclicity was measured as the minimum of positive and negative pulse peaks, counting all peaks with at least 2/3 the amplitude of the maximum peak within a pulse. A pulse was considered to be polycyclic when it had three or more peaks. For analysis of ipis, we evaluated only ipis between 15-80 ms in trains of more than 2 pulses. Carrier frequencies of pulse and sine song were assessed by plotting the median value of each fly.

Playback experiments were performed as described previously (Kerwin et al. 2020). Wings were amputated with spring scissors close to the hinge and flies were allowed to recover for at least 24 hrs before testing. The approximately 3 s long song examples from a *wt* and *fru>Gad1-IR* fly were continuously played back and had the following characteristics representative for their genotype: number of pulses; 38 and 34; % ipis between 30-40 ms: 86 and 41; % polycyclic pulses 3 and 41; median pulse frequency: 246 Hz and 322 Hz, respectively.

### Immunohistochemistry and confocal imaging

Immunohistochemistry and confocal imaging of fly nervous systems were performed as described previously (O’Sullivan et al. 2018). Fly nervous systems were dissected in PBS, fixed in 4% paraformaldehyde, stained with mouse nc82 antibody (Developmental Studies Hybridoma Bank, RRID: AB_2314866), rabbit anti-GFP antibody (Torrey Pines Biolabs, RRID: AB_10013661) or chicken anti-Gfp antibody (Abcam, RRID: AB_300789), rabbit anti-DsRed antibody (Living colors/Takara Bio, RRID: AB_10013483) and mounted in Vectashield (Vector labs, RRID: AB_2336789). Confocal image stacks were acquired at a Zeiss LSM780 microscope.

## Acknowledgements

We thank Asia Ahmed, Akhil John, Francesca Barbieri, Peter Kerwin and Yanan Zhang for help with experiments, Anna Prudnikova, Marketa Kaderakova and Tatiana Adamiec for technical assistance. We thank DGRC and VDRC stock libraries, Developmental Studies Hybridoma Bank for the nc82 antibody, Barry Dickson and Ben White for providing fly stocks. This study was supported by Lundbeckfonden grant DANDRITE-R248-2016-2518.

---

**Supplementary Figure 1.**
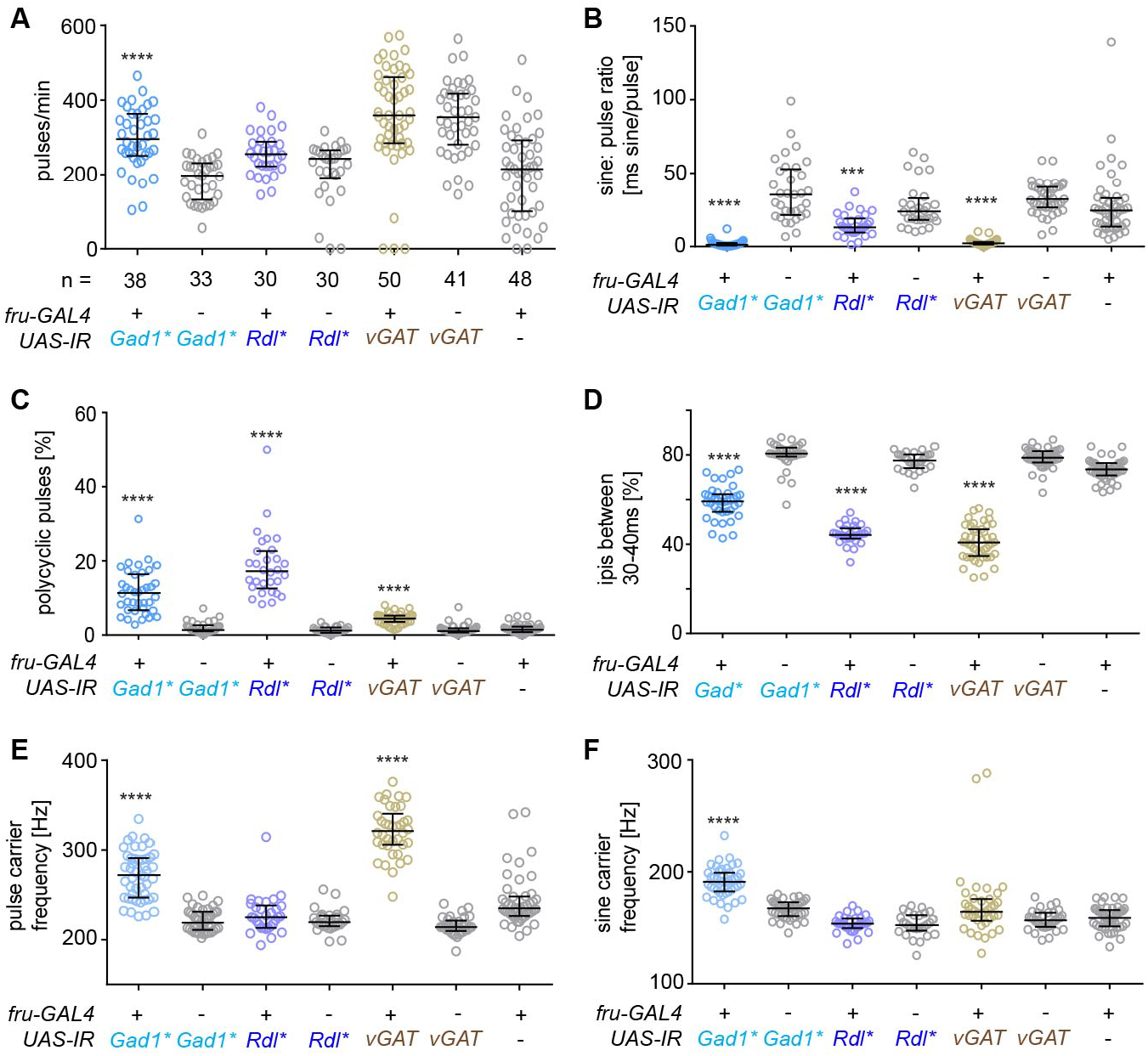
Song patterning defects upon knockdown of GABAergic signaling in *fruitless* neurons with additional RNAi lines. **A** Amount of courtship song (in pulses/min) displayed toward a virgin wild type female of male flies upon RNAi-mediated knockdown of *Gad1, Rdl* or *vGAT* with additional *UAS-IR* RNAi lines in *fruitless* neurons and respective genetic controls. n indicates number of flies tested. **B** Amount of sine song (sine: pulse ratio) for knockdown and control males. **C** Amount of polycyclic pulses (% of pulses with more than 2 cycles) in the song of knockdown and control males. **D** Percentage of inter pulse intervals (ipis) within the 30 – 40 ms range in the song of knockdown and Control males. **E** Median pulse song carrier frequency for knockdown and control males. **F** Median sine song Carrier frequency for knockdown and Control males. Each knockdown genotype is compared to its corresponding *UAS-IR* control and the *fru* ctrl, ***p = 0.0007, ****p< 0.0001, Kruskal-Wallis test with Dunn’s multiple comparison; n in **B-F** is the same as in **A**. All experiments are performed with wild type virgin females. In all scatterplots, each data point represent one fly, error bars indicate median with interquartile range.

## Notes

### Competing Interest Statement

The authors have declared no competing interest.

